# Guild-Level Gut Microbial Alterations Associate with Non-hepatitis Virus related Hepatocellular Carcinoma

**DOI:** 10.1101/2023.11.13.566887

**Authors:** Yubing Xu, Hongwei Ye, Xiaowei Meng, Shuyu Zhang, Yiyin Zhang, Yi Chen, Yu Chen

## Abstract

The gut microbiota of hepatocellular carcinoma (HCC) is different from that of cohort without HCC. However, the relationship between non-hepatitis virus related HCC and gut microbial alterations remains elusive. In this work, we studied the gut microbial data in Sequence Read Archive (SRA) database from 58 healthy controls (HC), 31 patients with nonalcoholic fatty liver disease (NAFLD), 206 patients with hepatitis B viruses, 35 patients with HBV related HCC (B-HCC), and 22 patients with non-HBV or non-HCV related HCC (NBNC-HCC). Depend on guild-based analysis, 27 gut bacterial co-abundance groups (CAGs) were established. By studying these CAGs, we found the gut microbial overall structure and composition differed among patients with NBNC-HCC, B-HCC, other liver diseases, and healthy cohort, such as members from Lachnospiraceae, Veillonellaceae, and *Prevotella*. The network calculated by these CAGs in patients with HCC showed less interactions than that in healthy cohort. The specific CAGs showed correlated to the development of HCC. In addition, gut bacterial dysbiosis index, which was calculated by significantly altered CAGs in patients with NBNC-HCC compared with healthy cohort, showed potential diagnostic power of NBNC-HCC. Therefore, this study shows a new way toward understanding the role of the gut microbiome in patients with non-hepatitis virus related HCC.

**IMPORTANCE:** Gut microbiome may play an important role in the pathogenesis and development of hepatocellular carcinoma. However, the relationship between non-hepatitis virus related hepatocellular carcinoma and gut microbiome should be further understood. In the current work, we found the alterations of gut microbial structure, composition, and network robustness in these patients by guild-based data analysis. Specific gut bacteria in these patients were related to clinical parameters, such as blood biochemical indexes. More importantly, gut bacterial bio-markers, which can discriminate these patients and cohort without this disease, were showed in this study. Thus, our study suggests that these guild-level gut bacteria may contribute to the development of non-hepatitis virus related hepatocellular carcinoma. This provides a new direction for future studies, which aims to show the importance of the gut microbiome in hepatocellular carcinoma.

---

Hepatocellular carcinoma (HCC), approximately 90% of liver malignancy cases, is the fifth cancer and the second leading cause of solid tumor related deaths worldwide (1). According to global epidemiological data, HCC has an incidence of more than 850,000 new cases annually (2, 3). The common risk factors of HCC are infection with hepatitis B and C viruses, nonalcoholic fatty liver disease (NAFLD), and ingestion of the fungal metabolite aflatoxin B1 (4). Approximately 80% cases of HCC are related to infections of HBV and HCV. However, the prevalence of non-HBV or non-HCV related HCC (NBNC-HCC) is elevating. Furthermore, 12-16% of all the HCC cases is NBNC-HCC according to epidemiological data based on Japanese cohort (5, 6). Compared to patients with HBV related HCC (B-HCC), patients with NBNC-HCC showed poorer prognosis, because NBNC-HCC was difficult to be detected early (7). Therefore, early detection of HCC, including B-HCC and NBNC-HCC, can significantly improve the prognosis (8, 9).

In the development of HCC, gut microbiome may play an important role. Gut microbial structure and composition in patients with HCC differed from those in healthy cohort (10). Furthermore, the microbial composition in NBNC-HCC patients altered compared with that in B-HCC patients (11). For instance, the relative abundance of Proteobacteria increased, while the Proteobacteria was decreased (12). Emerging evidence has demonstrated that alterations of gut microbiota affect liver diseases, such as NAFLD, the development of HBV infection, cirrhosis, and HCC (13, 14). Moreover, early patients with HCC showed enriched LPS-producing bacterial relative abundance, while decreased butyrate-producing genera (9, 15). The gut microbial translocation and metabolism changed the clinical outcomes of chronic liver diseases (16, 17). Gut microbiome can modulate the development and function of liver through the liver-gut microbiota axis, such as the regulation of synthesis of bile acids and butyric acid, and this may impact the development of HCC (18, 19). Zhang et al. found that high-fat/high-cholesterol diet induced NAFLD associated HCC by altering the gut microbial composition and metabolites in animal model. In addition, gut microbiota restored by the treatment of atorvastatin could prevent the development of NAFLD associated HCC, completely (20). Overall, these findings show that gut microbial alterations may contribute to the pathophysiology of HCC.

The reported findings analyzed the gut microbial alterations in HCC patients at low-resolution levels, including species, genus, or even phylum (9, 12). However, previous studies found that the bacterial functions were species- or even strain-specific (21-23). In the gut microbial ecosystem, the members whose function is similar show consistent co-abundant behavior and may work together to influence the pathophysiology and development of human diseases (24-28). Thus, the relationship between the gut microbiome and the development of NBNC-HCC merits further understood in an ecological way.

In this work, we analyzed the gut microbial data sets from previous studies (12, 29), including 58 healthy controls (HC), 31 NAFLD patients, 206 HBV patients, 35 B-HCC patients, and 22 NBNC-HCC patients. Compared with HC, NAFLD, and HBV patients, the gut microbial overall structure, composition, and relationship of network in patients with HCC were altered in guild-level. In HCC group, we found that specific gut bacteria were related to general anthropometric parameters and blood biochemical indexes. By guild-based data analysis, we found gut microbial bio-markers, which can discriminate among NBNC-HCC patients, with high sensitivity and specificity.

## RESULTS

### Gut microbial alterations by Guild-level data analysis in patients with HCC

We analyzed the gut bacterial 16S rRNA gene V3-V4 regions in fecal samples from Wang et al’s study (29) and V4 regions from Liu et al’s study (12). Liu et al’s cohort included 33 (HC), 35 B-HCC patients, and 22 NBNC-HCC patients. And Wang et al’s study recruited 25 healthy subjects, 31 NAFLD patients, 206 HBV patients.

Totally, 4,609,848 and 4,118,375 high-quality sequencing reads (51,220.5 ± 5,801.2 and 15,719.0 ± 3,010.5 reads per sample) were denoised into 2,082 and 4,279 amplicon sequence variants (ASVs) by QIIME2 in Liu and Wang et al’s study, respectively (30, 31). Then, we compared the representative reads between these studies. 990 representative reads (210 bp) of Liu et al’s study (12) can be matched completely in 2,554 (460 bp) of Wang et al’s study (29). Then, two gut microbial datasets were merged into a new matrix, which included 8,728,223 high-quality sequencing reads (24,796.1 ± 15,975.2 reads per sample), and these reads were denoised into 3,708 ASVs.

To explore the alterations in gut microbial compositions of HCC patients by an ecologically relevant way, we established a co-abundance network depend on Spearman’s correlation coefficients among the 354 ASVs that existed in 20% of volunteers in these 5 groups, and clustered these ASVs into 27 co-abundance groups (CAGs) for further analysis (Fig. 1). CAG (guild)-based analysis may provide a more ecologically relevant method to understand the relationship between gut microbiota and disease. In this method, we should consider both abundance and evenness of these CAGs, thus CAGi was calculated (32).

**FIG 1.**
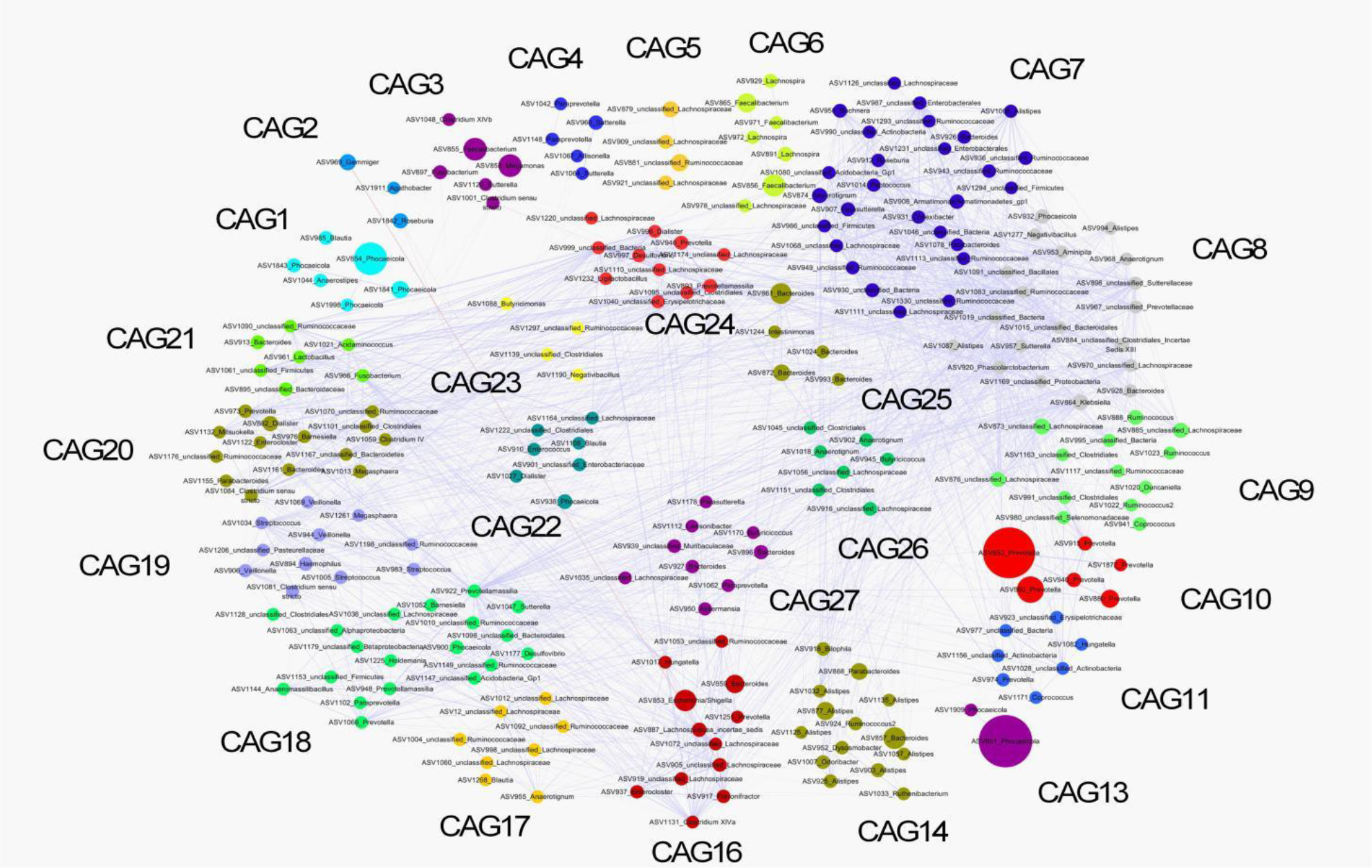
ASV-level network diagram of the 354 ASVs shared more than 20% of volunteers in the HC, NAFLD, HBV, B-HCC, and NBNC-HCC groups. Node size indicates the mean abundance of each ASV, with line width standing for correlation magnitude. Red lines and blue lines between nodes represent positive and negative correlations between the nodes which they connect, respectively. Only lines corresponding to correlations with a magnitude greater than 0.4 are drawn. The 354 ASVs are clustered into 27 gut bacterial CAGs based on their Spearman’s correlation coefficient. ASV, amplicon sequence variant; CAGs, co-abundance groups; HC, healthy controls (*n* = 58); NAFLD, Nonalcoholic fatty liver disease patients (*n* = 31); HBV, Hepatitis B virus infection patients (*n* = 206); B-HCC, HBV related hepatocellular carcinoma patients (*n* = 35); NBNC-HCC, non-HBV non-HCV related HCC patients, (*n* = 22).

Principal-coordinate analysis (PCoA) of the Jaccard and Bray-Curtis distance by the CAGi of CAGs showed that the overall structure of gut microbiota in the HCC group significantly differed from that in the HC, NAFLD, and HBV group (Fig. 2A, B, and Table. 1). In addition, we calculated the topological parameters of networks in HC, B-HCC, and NBNC-HCC to study the differences existed in complexity among the networks of CAGs. The total number of nodes among the 3 networks was same, however the total number of edges, network degree, and density, which is defined as the ratio of the number of actual edges and the number of possible edges, were decreased in HCC group compared with HC groups (Fig 3A, B, and C). Moreover, the network degree cumulative frequency, a measure of the relative connectivity of each node in a network, decreased significantly in HCC groups (Fig. 3D). These results suggested that HCC patients had less microbial interactions compared with healthy cohort.

**TABLE 1.** The perMANOVA of PCoA in the HC, NAFLD, HBV, B-HCC, and NBNC-HCC groups based on the 27 CAGs.

| Groups | *P* Value (Jaccard) | *P* Value (Bray Curtis) |
| --- | --- | --- |
| HC vs. B-HCC | 0.001 | 0.004 |
| HC vs. NBNC-HCC | 0.008 | 0.049 |
| NAFLD vs. B-HCC | 0.001 | 0.001 |
| NAFLD vs. NBNC-HCC | 0.001 | 0.001 |
| HBV vs. B-HCC | 0.001 | 0.001 |
| HBV vs. NBNC-HCC | 0.001 | 0.001 |
| B-HCC vs. NBNC-HCC | 0.966 | 0.255 |

**FIG 2.**
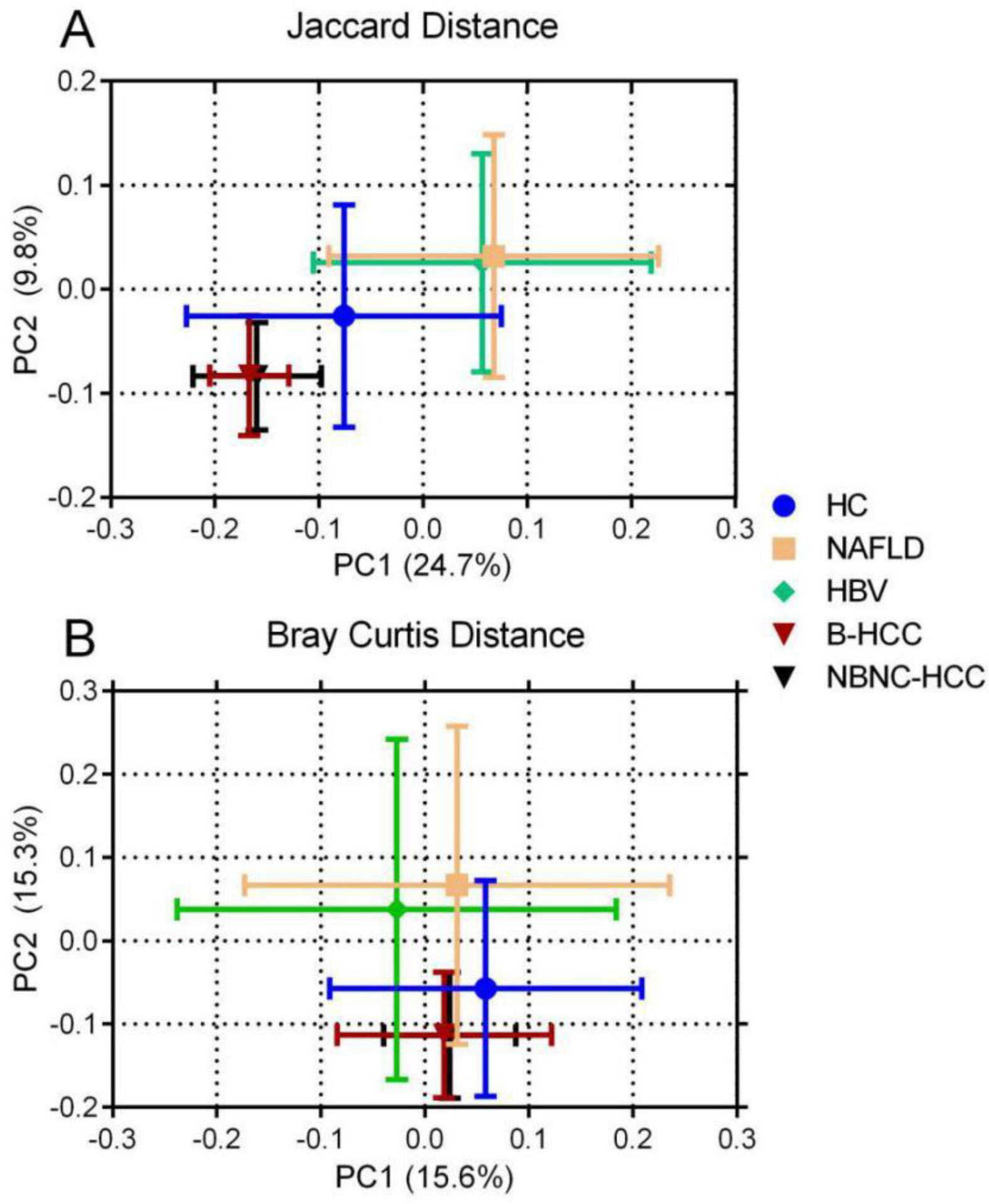
PCoA in the HC, NAFLD, HBV, B-HCC, and NBNC-HCC groups. (A) Jaccard distance. (B) Bray-Curtis distance. The circles indicate the mean of each group and the error bar represents the SD. HC, healthy controls (*n* = 58); NAFLD, Nonalcoholic fatty liver disease patients (*n* = 31); HBV, Hepatitis B virus infection patients (*n* = 206); B-HCC, HBV related hepatocellular carcinoma patients (*n* = 35); NBNC-HCC, non-HBV non-HCV related HCC patients, (*n* = 22).

**FIG 3.**
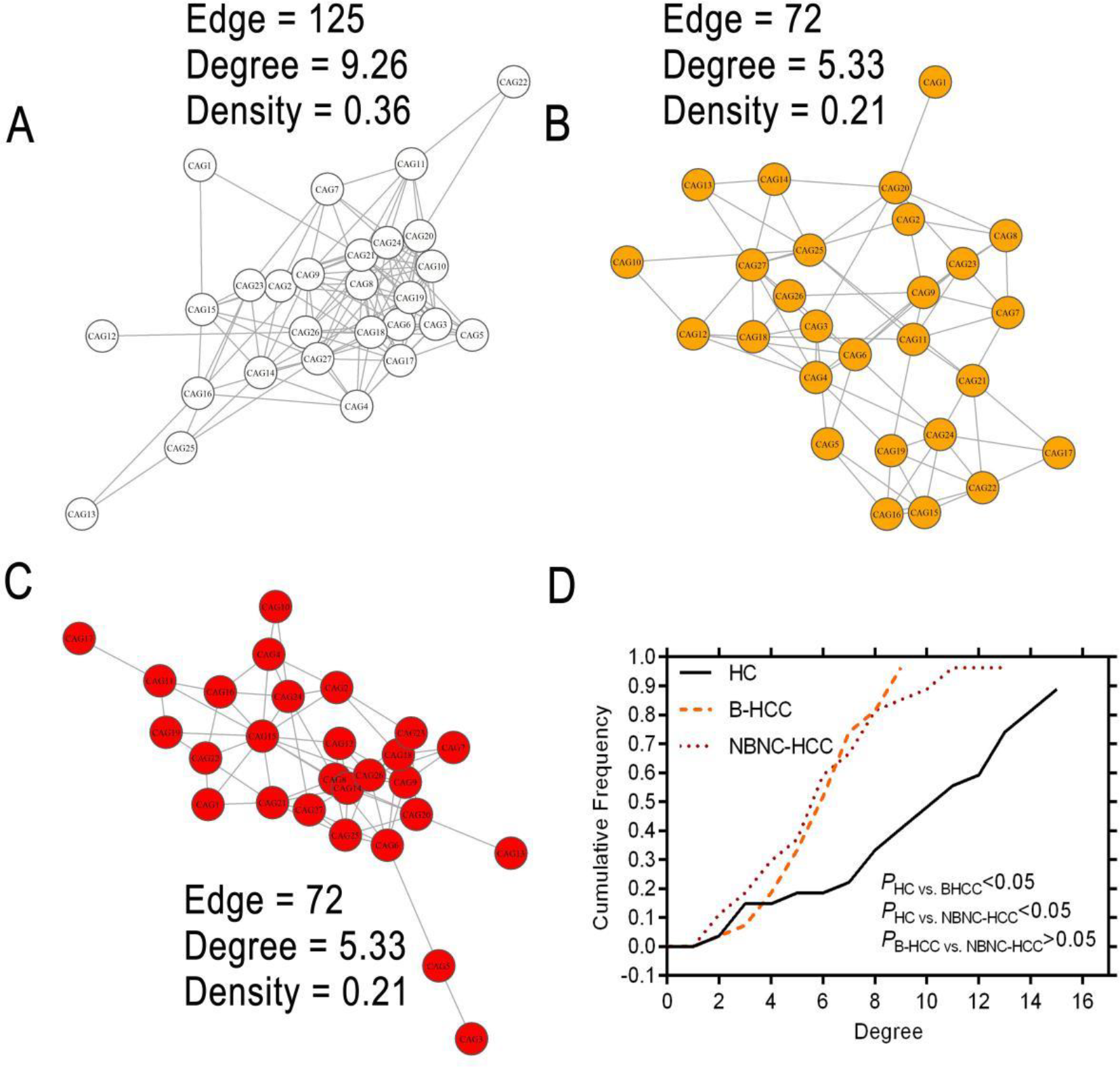
The ecological relationship of the gut microbial members. Visualization of constructed networks based on Spearman’s correlation coefficient from the HC (A), B-HCC (B), and NBNC-HCC (C) groups and degree centralities of networks form the 3 groups (D). The test of Wilcoxon rank sum was utilised for analyzing the variations between two groups in cumulative distributions. HC, healthy controls (*n* = 58); B-HCC, HBV related hepatocellular carcinoma patients (*n* = 35); NBNC-HCC, non-HBV non-HCV related HCC patients, (*n* = 22).

### Association of gut bacteria to clinical parameters in patients with HCC

The relationship of gut microbiome with general anthropometric parameters and blood biochemical indexes was studied by calculating Spearman’s correlation coefficients. The CAGi of CAG 16 was positively correlated to heigh, while CAG 11 showed significant negative correlation with heigh and weight in the HC, B-HCC, and NBNC-HCC groups (Fig. 4A). In HCC group, we found that CAG 3, 5, 8, 11, 13, 14, 17, 18, and 25 were positively related to blood biochemical indexes, however CAG 2, 3, 11, 15, 16, 20, 21, 22, and 24 showed significantly negative correlation (Fig. 4B).

**FIG 4.**
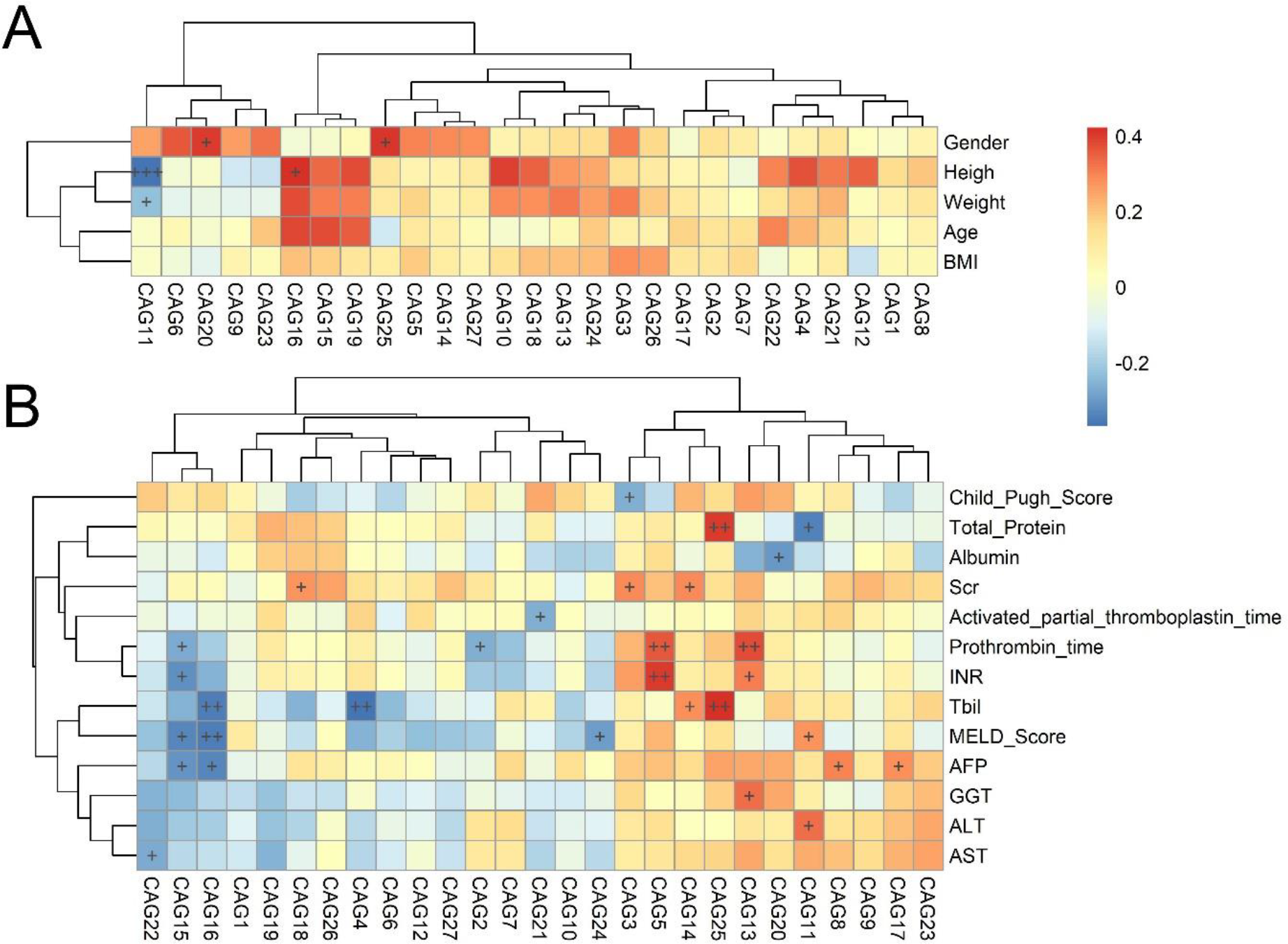
Heat maps of correlations between CAGs and clinical parameters in HCC patients. (A) General anthropometric parameters in the HCC and HC groups. (B) Blood biochemical indexes in the HCC group. The color of the cells represents Spearman’s correlation coefficient between each CAG and clinical parameter. \**P* < 0.05, \*\**P* < 0.01, \*\*\**P* < 0.001. BMI: body mass index, *n* = 90; AFP, alpha-fetoprotein, *n* = 54; ALT, alanine aminotransferase, *n* = 56; AST, aspartate transaminase *n* = 56; GGT, Gamma-Glutamyl Transferase, *n* = 56; TBil, total bilirubin, *n* = 56; INR, Prothrombin Time / International Normalization Ratio, *n* = 56; Scr, serum creatinine, *n* = 56.

### Specific gut microbial members distinguish the HCC groups with other groups

By statistical analysis, we found that CAGi of CAG7, 8, 10, 11, 19, 21, 22, and 24 was significantly increased in B-HCC group, while that of CAG8, 19, 22, and 24 was significantly increased in NBNC-HCC group (Fig. 5). CAG7, 8 mainly contained ASVs belonging to the family Ruminococcaceae, CAG10 to the genus *Prevotella*, CAG11 to the family Ruminococcaceae and Lachnospiraceae, CAG19 the family Veillonellaceae, CAG21 to the family Bacteroidaceae, CAG22 to the family Bacteroidaceae and Lachnospiraceae, and CAG24 to the family Prevotellaceae and Lachnospiraceae (Table S1).

**FIG 5.**
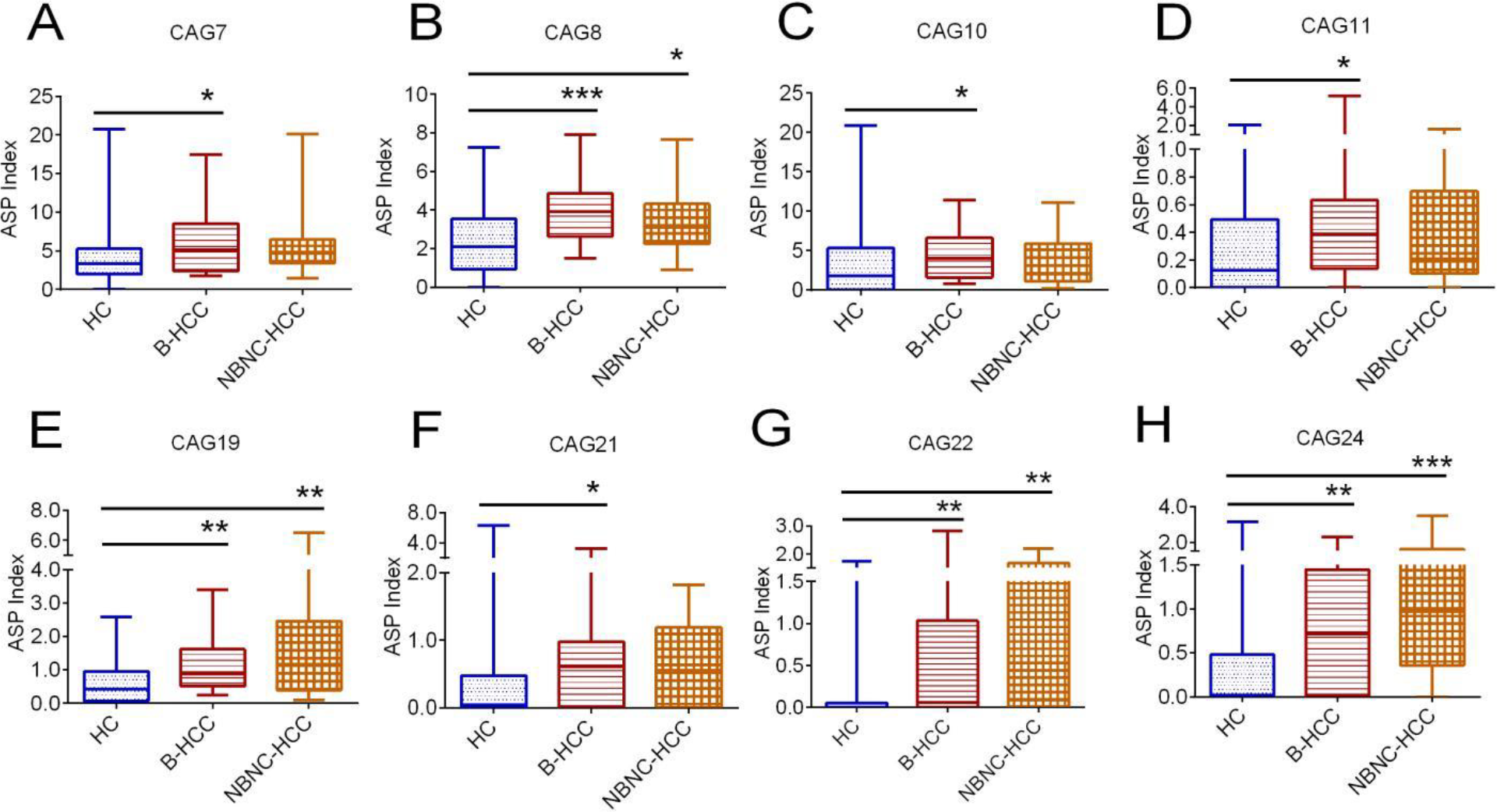
The gut bacterial alteration in the B-HCC and NBNC-HCC groups compared with the HC groups. (A) CAG7. (B) CAG8. (C) CAG10. (D) CAG11. (E) CAG19. (F) CAG21. (G) CAG22. (H) CAG24. The Kruskal-Wallis test was used for comparisons in groups. *, *P* < 0.05, **, *P* < 0.01, ***, *P* < 0.001. HC, healthy controls (*n* = 58); B-HCC, HBV related hepatocellular carcinoma patients (*n* = 35); NBNC-HCC, non-HBV non-HCV related HCC patients, (*n* = 22).

**FIG 6.**
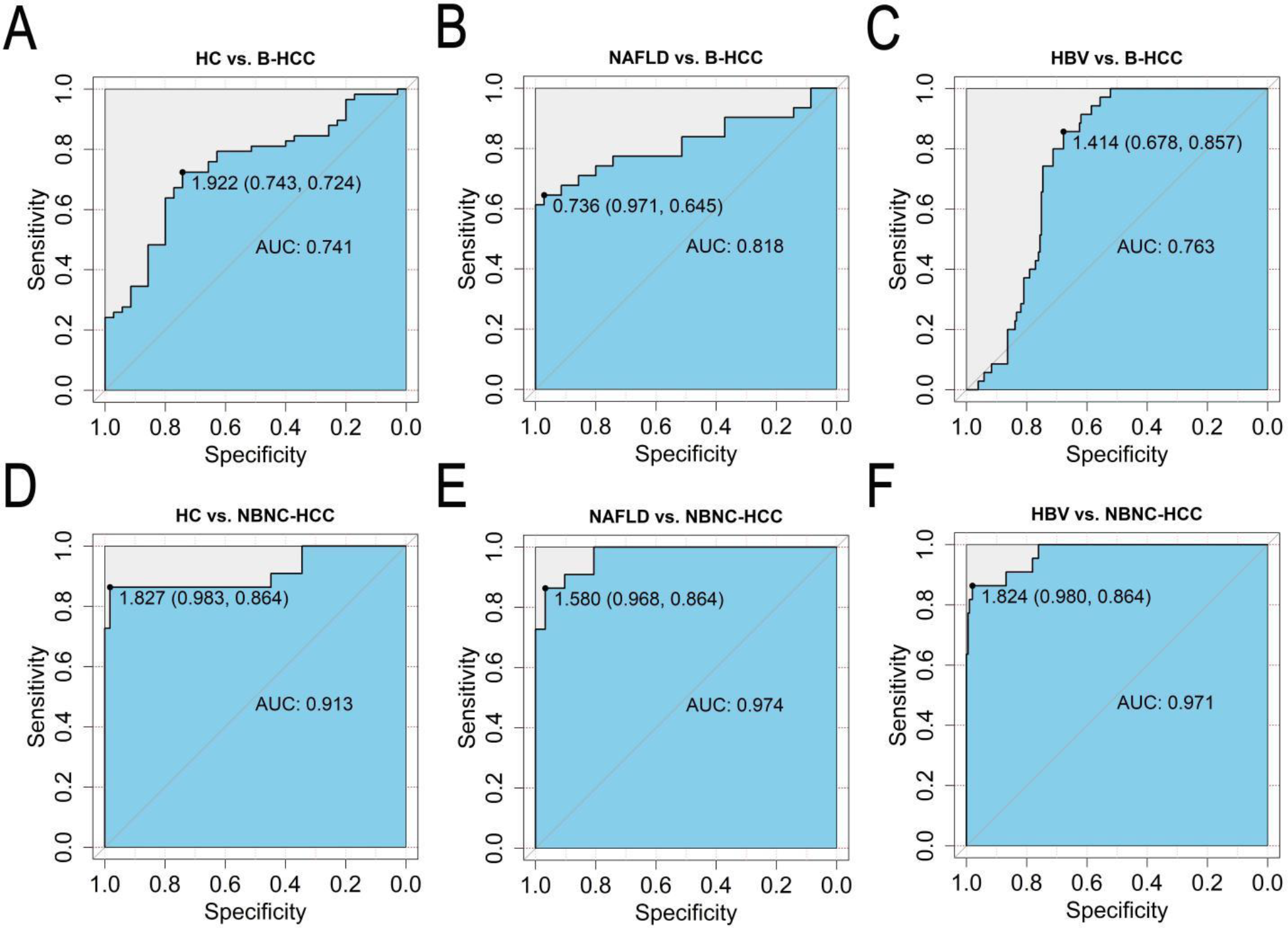
Diagnostic performance of fecal microbial markers for B-HCC and NBNC-HCC. ROC of HC versus B-HCC (A), NAFLD versus. B-HCC (B), HBV versus B-HCC (C), HC versus NBNC-HCC (D), NAFLD versus NBNC-HCC (E), and HBV versus NBNC-HCC (F). The point stands for the GBDI threshold chosen based on Youden’s J statistic, and the corresponding specificity and sensitivity.

We calculated the gut bacterial dysbiosis index of B-HCC (B-GBDI) and NBNC-HCC (NBNC-GBDI) based on the significantly differential CAGs in HC vs. B-HCC and HC vs. NBNC-HCC, respectively. Receiver operating characteristic (ROC) curves were used to define a diagnostic threshold of HCC based on the GBDI values. The area under the ROC curves (AUC) of 0.913, 0.974, and 0971 showed promising potential diagnostic power of NB-GBDI in HC vs. NBNC-HCC (Fig 5D), NAFLD vs. NBNC-HCC (Fig 5E), and HBV vs. NBNC-HCC (Fig 5F). However, we didn’t find powerful predictive performance of B-GBDI (Fig 5A, B, and C). We chose the optimal NBNC-HCC threshold at the cut-off value which maximized the Youden’s J statistic. The NBNC-HCC threshold between HC and NBNC-HCC was 1.827, and corresponding sensitivity and specificity were 0.983 and 0.864.

## DISCUSSION

In the current study, we found altered gut microbiome in patients with HCC compared with healthy cohort and other liver disease, such as NAFLD and HBV infection. The gut microbiota in patients with HCC showed less robustly relationship than those in healthy cohort. Furthermore, we found specific gut bacteria, such as the members from the family Ruminococcaceae, Lachnospiraceae, Bacteroidaceae, Prevotellaceae and the genus Prevotella, were correlated to the development of HCC and acted as potential non-invasive bio-markers for earlier and auxiliary diagnosing non-hepatitis virus related HCC.

In our work, the gut microbial overall structure and network’s topology were changed in patients of HCC compared with patients with HBV infection and healthy cohort, which is consistent with a previous study based on Chinese cohort (9, 10). Moreover, the gut microbial composition in patients with HCC was different to that of healthy cohort. For instance, the specific gut microbial members, which were mainly from the family Ruminococcaceae, Bacteroidaceae, Lachnospiraceae, Veillonellaceae and the genus *Prevotella*, increased in patients with B-HCC. Previous research article found that the relative abundances of these families or genus were enriched in patients with HCC, which is consistent with our study (33). However, a previous study revealed that Ruminococcus and Lachnospiraceae were significantly decreased in Chinese patients with HCC, which is different from our study. On the other hand, even in our research, the specific gut bacterial ASVs from Ruminococcus and *Prevotella* didn’t significantly alter in patients with HCC compared with healthy cohort (Table S1). These results suggested that the alterations in these microbial features belonging to the same family or genus may be different between patients with HCC and healthy cohort. The phenomenon might be attributed to the reason that the bacterial functions are strain-specific (34, 35). Therefore, future studies should explore the relationship between the gut microbiota and NBNC-HCC at the strain level.

Because of the strain-level bacterial functions in our gut, microbial data sets show highly dimensional and sparse characteristics (25, 28). Guild-based analysis could reduce the dimensionality of gut microbial datasets in an ecologically relevant way compared with the conventional method, such as taxon-based analysis, to identify the critical members of the gut microbiota related to human disease (28). Members in a same guild may show co-abundant behavior and work together to influence human health and diseases (24). In our work, we found specific gut bacteria, such as members from the family Ruminococcaceae, Bacteroidaceae, Lachnospiraceae, Veillonellaceae and the genus *Prevotella*, elevated in patients with HCC by this type of analysis. The gut-liver axis, primarily through the portal circulation, is a bidirectional interaction of the liver and gastrointestinal tract. The relationship between the gut microbiome and the liver is regulated by a complex network of interactions, such as metabolize, immunity, and neuroendocrine (36, 37). Alterations in these gut bacteria have been reported to play a role in the abnormalities of gut-liver axis, which may contribute to the development of HCC (20, 38, 39). Many members of the family Lachnospiraceae are regarded as lipopolysaccharides (LPS)-producing bacteria (29). LPS activates NF-κB pathway, which produces pro-inflammatory cytokines, such as TNF-α and IL-6 (40). This leads to liver inflammation and oxidative damage, then promoting the occurrence and development of HCC (20). Specific bacteria from the family Veillonellaceae increase intestinal permeability, which enable more microbial metabolites to reach the liver, then may influence the metabolism of bile acid to impair the function of liver (41, 42). Thus, our work identified these bacteria, which altered in patients with HCC, may contribute to the pathophysiology and development of this disease. Next, we should understand the mechanisms by which these gut bacteria contribute the symptoms of NBNC-HCC.

The clinical feature profiling of patients with NBNC-HCC is different from that of patients with B-HCC. Compared with patients with B-HCC, more patients with NBNC-HCC are detected at an advanced stage (42). For earlier diagnosis, it is important to find non-invasive bio-markers to indicate the development of NBNC-HCC. In the current study, we find an index, which are calculated by the richness and diversity of specific gut bacteria, can potently distinguish between patients with NBNC-HCC and healthy cohort, even between NBNC-HCC and HBV, NBNC-HCC or NAFLD. In the future, prospective clinical research with a larger cohort should be performed to validate the potential diagnostic power of this index.

In conclusion, our study demonstrates the alterations of gut microbiota in patients with HCC. In addition, guild-level gut microbial signature can distinguish NBNC-HCC from healthy cohort and other liver disease. Specific gut bacteria may play an important role in the development of NBNC-HCC, providing a new direction for future studies aiming to explore the role of the gut microbiome in NBNC-HCC.

## MATERIALS AND METHODS

### Clinical Investigation

All participants in this clinical trial were from Liu et al.’s (12) and Wang et al.’s (29) studies. Liu et al.’s study comprised 33 healthy participants, 35 patients with B-HCC, and 22 patients with NBNC-HCC, while Wang et al.’s comprised 33 healthy participants, 31 patients with NAFLD, and 206 patients with HBV. The clinical parameters, such as inclusion criteria, exclusion criteria, pathologic diagnosis, the results of blood test, were from these two studies.

### Bioinformatics analysis of Amplicon sequencing data

Amplicon sequence data of gut microbial 16S rRNA gene V3-V4 (12) and V4 (29) region from the Illumina Miseq platform was downloaded from the NCBI Short Read Archive (SRA) under BioProject PRJNA382861 and GSE108847, respectively. The data were imputed into Quantitative Insights Into Microbial Ecology 2 (QIIME2), and the steps of adapter sequences removing, DADA2 pipeline (error filtering, trimming, denoising, merging of paired reads, and removal of chimeras), and amplicon sequence variants (ASVs) clustering were performed in this platform (version 2021.11). We compared the representative reads from Liu et al.’s (12) study with those from Wang et al.’s study (29). The reads from Liu et al.’s study (12) that could be matched on 150-460 bp region of reads from Wang et al.’s study (29) was considered as the same and analyzed in next step. All ASVs were annotation by Ribosomal Database Project (RDP). The reads of each sample were downsized to 10,000 to normalize the even sampling depths.

### Network construction

The relationships of correlation among 354 ASVs, which colonized in more than 20% of volunteers in these groups, were calculated by the Spearman’s correlation coefficient (R) depend on their relative abundance. This R matrix was converted to a correlation distance (1-R) matrix by WGCNA package in the R (version 4.1.2). Based on the Ward.D2 clustering algorithm in this package, a hierarchical clustering tree of the distance matrix was established, then these ASVs were clustered into 27 CAGs. By using Cytoscape (version 3.2.1), the network of these ASVs was visualized. We proposed a CAGi to measure the abundance and evenness of CAG which is expressed by equation below:

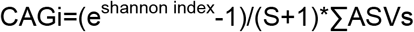

S is the number of ASVs in each CAG and ΣASVs is the summed abundance (32).

By utilizing Spearman’s correlation coefficient as weight, layout of nodes and edges was calculated by the Fruchterman-Reingold. The calculation and visualization of networks’ topological characteristics were showed by igraph package in the R (version 4.1.2).

### Definition of the NB/B-GBDI

NB/B-GBDI was considered as indexes to evaluate the severity of the gut microbial CAGs alterations in patients with NBNC/B-HCC compared with healthy cohorts. The equation was showed below:

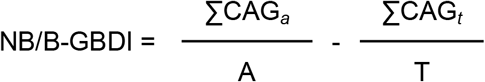

CAG_*a*_ represented the NBNC/B-HCC patient-enriched CAGs, while CAG_*t*_ represented the HC-enriched. A and T represented the number of CAGs belonging to CAG_*a*_ and CAG_*t*_, respectively. If A of a CAG is 0, we defined ΣCAG_*a*_/A is equal to 0, and the same as ΣCAG_*t*_/T.

### Statistical analysis

Permutational multivariate analysis of variance (perMANOVA) was utilized to calculate the significant difference of CAGs overall structure among groups by “vegan” package in the R (version 4.1.2). Kolmogorov-Smirnov test was utilized to estimate the topological characteristics of CAGs network among these groups. Spearman’s correlation coefficient was used to compare the correlation between CAGs and the clinical parameters of volunteers. Kruskal-Wallis test was performed to analyze the significant difference of CAGi among these groups by using the statistical software SPSS26.0 (IBM Corp., Armonk, N.Y., USA). ROC was also calculated by SPSS26.0. *P* values < 0.05 were known as statistically significantly difference.

## Supplementary Material

Table S1 Taxonomical assignments of ASVs in 27 CAGs.

ASVs: Amplicon Sequence Variants; CAG: Co-abundance group

